# Effect of elevated embryonic incubation temperature on the temperature preference of juvenile lake (*Coregonus clupeaformis*) and round whitefish (*Prosopium cylindraceum*)

**DOI:** 10.1101/2023.03.13.532459

**Authors:** Adam A. Harman, Hannah Mahoney, William Andrew Thompson, Meghan L.M. Fuzzen, Bhuvan Aggarwhal, Lisa Laframboise, Douglas R. Boreham, Richard G. Manzon, Christopher M. Somers, Joanna Y. Wilson

## Abstract

Anthropogenic impacts can lead to increased temperatures in freshwater environments through thermal effluent and climate change. Thermal preference of aquatic organisms can be modulated by abiotic and biotic factors including environmental temperature. Whether increased temperature during embryogenesis can lead to long-term alterations in thermal preference has not been explicitly tested in native freshwater species. Lake (*Coregonus clupeaformis*) and round (*Prosopium cylindraceum*) whitefish were incubated at natural and elevated temperatures until hatching, following which, all groups were moved to common garden conditions (15°C) during the post-hatching stage. Temperature preference was determined at 8 (Lake whitefish only) and 12-months of age (both species), using a shuttlebox system. Round whitefish preferred a cooler temperature when incubated at 2°C and 6°C compared to 0.5°C. Lake whitefish had similar temperature preferences regardless of age, weight, and incubation temperature. These results reveal that temperature preference in freshwater fish can be programmed during early development, and that round whitefish may be more sensitive to incubation temperature. This study highlights the effects that small increases in temperature caused by anthropogenic impacts may have on cold-adapted freshwater fish.

## Introduction

Anthropogenic impacts have created scenarios where animals may be experiencing thermal stress during early critical life stages. Current predictions have proposed that the great lakes are expected to rise in temperature between 4-6°C by 2100 (IPCC, 2014), at a rate of about 0.1°C/year (Austin and Colman, 2007). More immediate concerns arise from warmer effluent discharge from industrial practices that use natural bodies of water to remove waste heat. This effluent can lead to increases in local habitats by 1-3°C, particularly those along the shoreline (Thome et al., 2016). Nearshore environments are critical regions for aquatic species, such as fish, providing regions to forage, shelter, and breed (Hampton et al., 2011). One risk for aquatic species that breed in these environments is the exposure of immobile embryos to supraoptimal temperature conditions throughout embryogenesis.

Environmental temperature exerts considerable control over the chemical process orchestrating development in fish (Stevens and Fry, 1970). Water temperature is a key determinant of growth in fish (Jobling, 1981; Magnuson et al., 1979), with increased temperature during embryogenesis leading to accelerated growth and developmental rates (Gillooly and Dodson, 2000; Nytrø et al., 2014; Sun and Chen, 2014). However, fish embryos display many plastic traits that can be influenced by their developmental environment (Jonsson and Jonsson, 2019). For instance, a positive relationship has been noted with incubation temperature and post-hatch metabolic rate in several fish species (Barrionuevo and Burggren, 1999; Bozek et al., 1990; Marty et al., 1990). Interestingly, the thermal optimum of key metabolic enzymes at the adult stage increases in response to temperatures experienced during rearing (Schnurr et al., 2014). Taken together, this may be indicative of an increased need to elevate body temperature to meet changes in metabolic demands. As ectothermic poikilotherms, several studies have demonstrated that fish aggregate to their thermal preference (T_pref_; Kellogg and Gift, 1983; Reynolds and Casterlin, 1980; Stevens and Fry, 1970), to maintain their metabolic, growth, and/or reproductive optimums (Haesemeyer, 2020; Larsson, 2005). T_pref_ has been shown to vary across life-stage (Edsall, 1999), season (Mortensen et al., 2007), time of day (Macnaughton et al., 2018) and metabolic state (Killen, 2014). However, the impact of elevated incubation temperature and any long-term change to T_Pref_, particularly in native cold-water fish, has not been explicitly tested.

Lake (*Coregonus clupeaformis*; LWF) and round (*Prosopium cylindraceum*; RWF) whitefish are cold-water adapted species that have an extensive range across North America (Ebener et al., 2010). These fish species serve an important ecological role in food webs and support commercial fisheries and indigenous communities (Ebener et al., 2010). LWF and RWF occur sympatrically, co-existing due to differential habitat and resource usage within lakes (Eberts et al., 2016). Both species of whitefish broadcast spawn in shallow (<10m) cobble beds in late November, and embryos remain in these shallow waters until the ice melt in spring (April-May; Scott and Crossman, 1973). These long incubation periods coincide with temperatures of 0.5-2°C at these depths (Patrick et al., 2013; Schwab et al., 1999; Thome et al., 2016), and these animals may be sensitive to increases in temperature imposed by anthropogenic impacts. Indeed, laboratory studies strongly support this, with whitefish exposed to elevated temperature during embryogenesis generally exhibiting perturbed morphology, precocious development, and increased mortality (Brooke, 1975; Eme et al., 2015; Eme et al., 2018; Lee et al., 2016; Lim et al., 2017; Lim et al., 2018; Mitz et al., 2019; Mueller et al., 2015; Mueller et al., 2017; Patrick et al., 2013; Price, 1940). These effects become more prevalent at constant temperatures ≥5° (Price, 1940; Brooke, 1975; Lim et al., 2017; Lim et al., 2018; Mitz et al., 2019; Eme et al., 2015; Eme et al., 2018; Mueller et al., 2015; Mueller et al., 2017; Patrick et al., 2013), with RWF appearing more sensitive, experiencing mortality rates 30-40% higher than LWF (Lim et al., 2017; Lim et al., 2018). Thermal stress during embryogenesis can augment and perturb the typical development of both LWF and RWF, but studies exploring impacts at post-hatch stages are limited. Work at later life-stages is a necessity given that whitefish embryos could be exposed to temperatures as high as 5°C now and up to 8°C within ~30 years at the current rate of warming (Austin & Colman, 2007).

This study tested the hypothesis that elevated temperature during rearing could impact the resulting thermal preference of juvenile LWF and RWF. Elevated temperature can lead to lethality in embryos of these species, but sublethal effects, such as changes in length and weight (Brooke, 1975; Price, 1940; Mitz et al., 2019; Lee et al., 2016), may lead to altered performance and function at later life-stages. To test this, we reared LWF at their optimum and natural rearing conditions (2°C) and elevated constant water temperatures of 5 and 8°C. As RWF are more sensitive to elevated incubation temperature and experience nearly 100% mortality at 8°C (Lim et al., 2017; Lim et al., 2018), we exposed RWF to 2°C and 6°C, and a colder temperature (0.5°C), to see effects in a lower range of environmental temperatures. We assessed behavioral performance at 8 and 12 months for LWF, and 12 months for RWF, determining their T_Pref_, velocity, total distance travelled, and movement across temperature gradients. The results suggest that elevated incubation temperature can alter RWF T_Pref_, but not LWF.

## Methods

### Study Species

Fertilized LWF embryos were acquired from Sharbot Lake White Fish Culture Station (Sharbot Lake, ON) on November 30, 2017 (reared to 12-months) or November 27, 2018 (reared to 8-months). Spawning RWF were obtained from Lake Ontario (Port Darlington, GPS 43°51’50”N 78°44’35”W) on December 10 and 11, 2018. RWF were stripped of eggs and milt and returned to the water. Artificial in-vitro fertilization occurred immediately after stripping. Embryos were disinfected with Ovadine^®^ solution and transported in lake water back to McMaster University. Embryos (160-310) were plated into 200mm x 20mm sterile petri dishes containing 200mL of dechlorinated city tap water, and then moved to incubators (Mitz et al., 2014). Embryos were initially kept at 8°C and cooled (1°C/week) until they reached a base temperature of 2°C, 5°C or 8°C for LWF or 2°C or 6°C for RWF (Fig. 1). To create the treatment for the 0.5°C RWF embryos, an ice slurry was maintained within an incubator set to 2°C. Incubation temperature was maintained for 100 days to replicate the winter period (December to March), following which, embryos were warmed (1°C/week) until reaching 8°C. To confirm temperature within each incubator, TidbiT^®^ temperature loggers placed in 200mm x 200mm petri dishes with 200 mL of dechlorinated water. For the base temperature (excluding warming and cooling periods), LWF were exposed to 2.08 ± 0.3°C, 4.81 ± 0.3°C, 8.05 ± 0.1°C, and RWF to 0.54 ± 0.2°C, 2.58 ± 0.2°C, 6.13 ± 0.2°C. Median hatch for LWF occurred at 50 days post fertilization (8°C), 108 days post fertilization (5°C), and 158 days post fertilization (2°C). Median hatch for RWF occurred at 88 days post fertilization (6°C), 114 days post fertilization (2°C), and 118 days post fertilization (0.5°C). Hatchlings (~10) were placed in 100mm x 20mm petri dishes with 100 mL of water at 8°C until successful exogenous feeding. Water in petri dishes was changed three times a week for embryos and daily for larvae. Larvae were transferred to 1-10L recirculating tanks and warmed (1°C/week) to 15°C, where they remained until testing (8- or 12-months post hatch). All treatment groups were maintained in common garden conditions once they were warmed to 15°C. Larval fish were initially fed *Artemia* nauplii and slowly transitioned to pellet feed (Otohime B1 (200-360 μm) – C2 (920-1,410 μm) larval feed).

**Figure 1.**
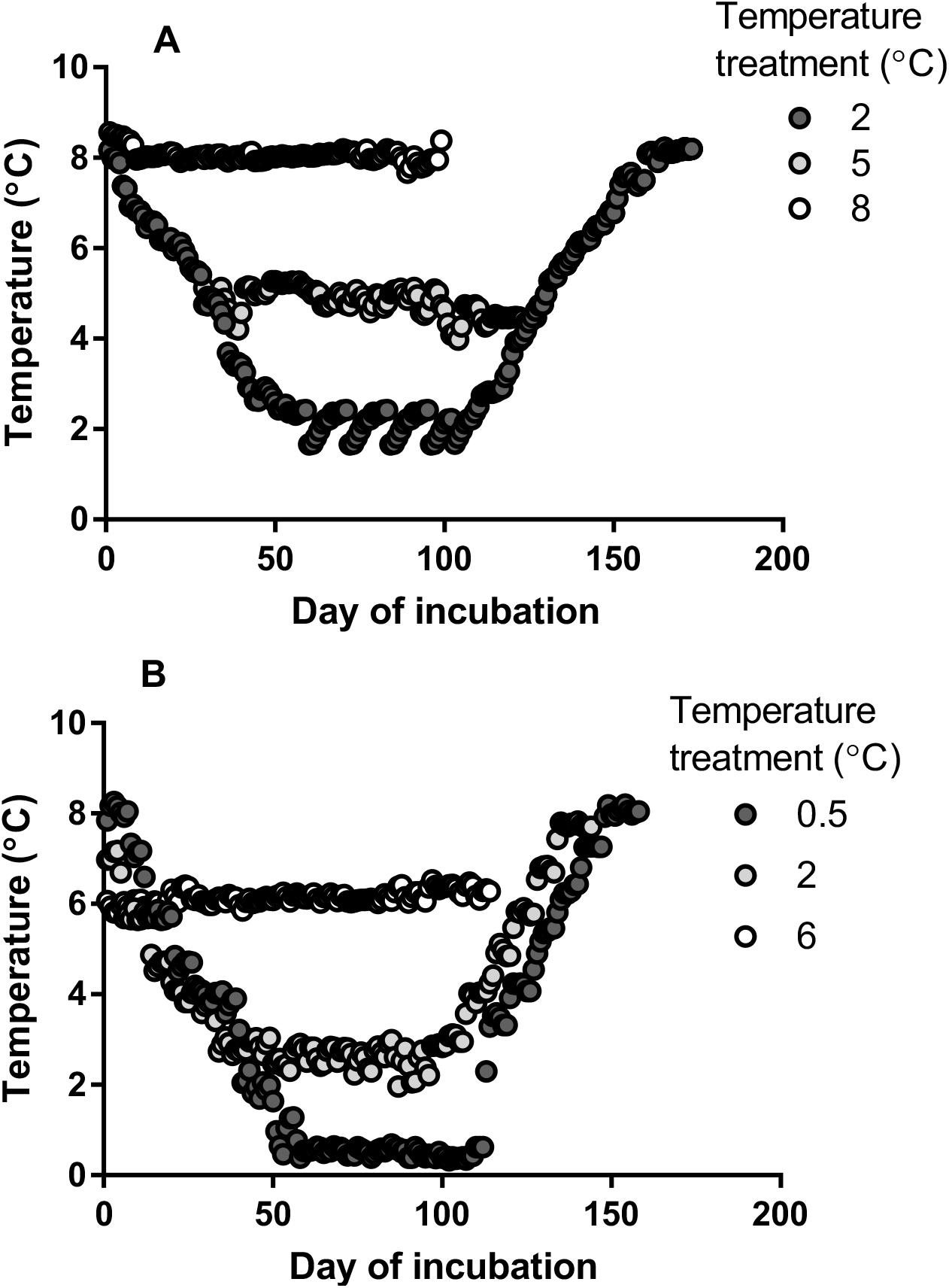
Daily temperature readings of incubation of lake whitefish (LWF; A) and round whitefish (RWF; B) until complete hatch of population. Data points represent averages of daily readings taken in 15 min intervals, with error bars excluded for visibility.

### Behavioral Assay – Shuttle Box

The shuttle box system (Loligo^®^), first described by Neill et al. (1972), consists of two cylindrical tanks connected by a small rectangular ‘shuttle’ to allow movement of animals between the tanks. Each tank is designated as the increasing (INCR) or decreasing (DECR) side, indicating the direction of temperature change when fish occupy that tank. To regulate temperature, system water was pumped through heat-exchange coils in hot (28°C) and cold (4°C) water baths (60L chest coolers) with mixing in separate buffer tanks for each side. A Recirculator 1/4 HP Chiller, Magnetic Drive Centrifugal Pump (300W/600W/950W @ 0°C/10°C/20°C; VWR) and 2×400W aquarium heaters were used to maintain the temperatures in the cold and warm bath, respectively. The shuttle box temperature probe can report temperature units to 0.01°C accuracy. Polystyrene insulation (1/2”) and foam insulation tape (1/4”) were used to prevent heat loss and maintain stable temperatures in the cold-water bath. System water flowed at 240 mL/min via gravitational pull through temperature probes and into the shuttle box where mixing between the two sides is minimized by counter-directional currents. The orientation of the INCR and DECR tanks and the side to which the fish would be introduced were randomized for each individual, using an online tool (random.org), to limit any potential bias introduced by visual cues or side preference. Whitefish of the appropriate treatment group were randomly selected from their home tank (15°C) and transported to the shuttle box system in blacked-out 1L glass beakers to prevent undue stress. A plastic divider separated the two halves of the arena, which when removed, started the acclimation period. Fish were acclimated to the arena in a static setting, with the two arenas maintained to 14 and 16°C with a hysteresis of 0.25°C. After 2h in this condition, the fish were tracked using a USB 2.0 uEye Camera tracked juvenile fish under infrared light (Loligo^®^ Infrared Light Tray), recording the position of the fish in the arena. The onset of warming or cooling occurred in response to whether the fish would be in the INCR or DECR tank, with the difference in temperature between these two sections being maintained at 2°C and warming or cooling (hysteresis = 0.1°C) occurring at a rate of 4°C/hour.

A maximum temperature of 23°C and a minimum temperature of 7°C was implemented to prevent exposure to extreme temperatures, which could cause stress or mortality (Edsall and Rottiers, 1976). T_Pref_ was calculated by the software as the median occupied temperature; additional measurements calculated were velocity (cm/s), total distance travelled (cm), time spent in INCR/DECR, number of passages, and avoidance temperature (temperature at which a passage between tanks occurred). Following the completion of the assay period, fish were removed, and measured for total length and mass before returning to a separate home tank (15°C). Prior to experimentation, whitefish were fasted for 12-20 hours to prevent fouling of the water and to standardize metabolic state. To account for any potential growth over the study duration, the order of sampling among treatment groups was randomized using an online tool (random.org). In total, 103 (12-month-old) and 87 (8-month-old) LWF, and 83 (12-month-old) RWF were tested for T_pref_ using the Loligo^®^ shuttle box system. Differences in treatment group sizes were due to differential mortality in holding tanks during rearing and were not due to experimentation. Prior to experimentation, power analyses were carried out to determine the optimal sample size within an acceptable power range (0.6-0.8; Harman et al., 2021).

### Statistical Analysis

Data is presented as mean±SD unless otherwise stated. T_Pref_, velocity, total distance travelled, time in arena, number of passages, and avoidance temperature between groups were analyzed using one-way ANOVA, with Tukey’s HSD post-hoc for comparisons between individual groups. To determine whether length or body weight influenced recorded T_Pref_, a general linear model was performed. A comparison was performed to assess species specific differences, with the 12-month T_Pref_ of RWF and LWF compared using a two-tailed T-Test. Bonferroni correction was applied to correct for multiple comparisons of T_Pref_, resulting in an α taken of 0.00625 for T_Pref_ analyses with LWF (a total of 4 comparisons), and 0.025 for RWF comparisons (a total of 2 comparisons). We developed a relationship between time (s) and temperature change (°C) in the shuttle box to determine the upper threshold of the system. This was done to remove possible outliers, as certain individuals were too active for the shuttle box system to determine T_Pref_ due to limitations in heating/cooling rates. Outliers were identified using the robust regression and outlier removal method (ROUT; Motulsky and Brown, 2006). The residuals of this fit were analyzed for potential outliers, and then subjected to ordinary least-squares regression after the removal of outliers. A total of 4 outliers were identified and removed using the ROUT method (2 x 8-month-old LWF, 2 x 12-month-old RWF). All statistical analyses were completed in R (version 4.0.0 “Arbor Day”), except for outlier identification which was completed in Graphpad Prism (version 8.4.3). All data and R scripts used for analysis were uploaded to a public GitHub data repository (https://github.com/WilsonToxLab/Shuttlebox-Thermal-Preference).

## Results

There was not a significant effect of rearing temperature on T_Pref_ on LWF of 8 months of age (Fig. 2A; F _[2,82]_ = 3.505; p=0.0346). Upper and lower avoidance temperatures were similar between treatment groups (Table 1). However, we note a non-significant trend, with eight-month-old LWF in the 5°C treatment group displayed the lowest activity, travelling an average distance of 173 m, compared to just over 190m at 2°C and 8°C. There was no observable change in the number of transitions between arenas in the shuttlebox in these fish. (Table 1). Total body length was similar between all treatment groups, varying less than 1mm on average (Table 1). Likewise, body weight was similar across treatment groups, with the largest difference (11%) between 2°C (1.13 ± 0.32g) and 5°C (1.25 ± 0.39g). Linear models were fit, including body weight and total length as fixed effects, to determine if there was a relationship between size and T_pref_. Model results (p = 0.068, p = 0.061) indicated there was no significant interaction between T_pref_ with total length or body weight.

**Figure 2.**
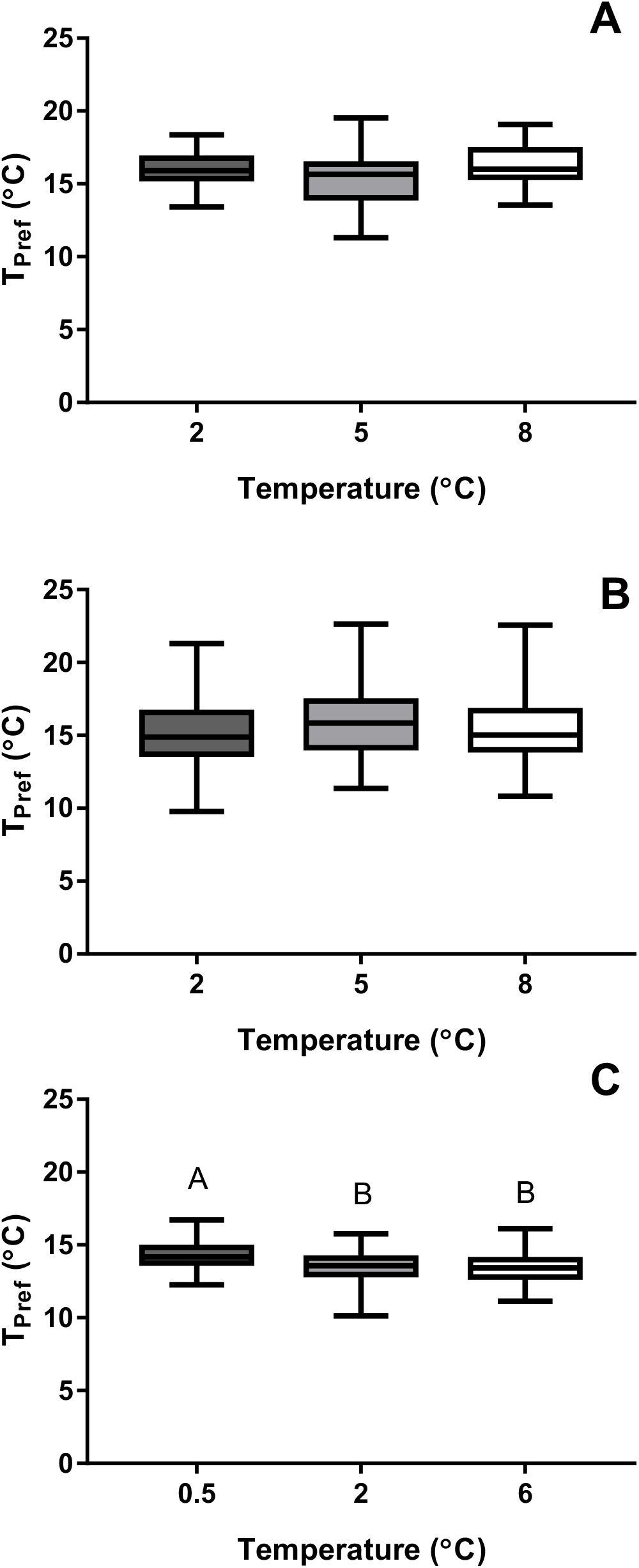
Ambient temperature during rearing can alter the thermal preference of fish in a species-specific manner. Boxplots comparing thermal preference (T_Pref_) of (A) 8-month-old lake whitefish, (B) 12-month-old lake whitefish, and (C) 12-month-old round whitefish, after exposure to different ambient temperatures during embryogenesis. Lower and upper box boundaries are 25^th^ and 75^th^ percentiles of the data, with the line inside the box representing the median, and error lines encompassing the entirety of the spread of data. Different letters above the boxplots denote significant differences between treatment groups.

**Table 1.** Behavioral output and whitefish characteristics of thermal preference experiment. Fish were reared at different temperatures during embryogenesis, and then held in common garden conditions post-hatch. Lake whitefish and round whitefish embryos were brought in at 8°C and cooled at 1°C/week to the base temperature (incubation treatment); after 100 days incubation, temperature was warmed at 1°C/ week until they reached 8°C. This simulated natural conditions with different base incubation temperatures. After successful exogenous feeding, fish were warmed at 1°C/week to 15°C and were held at that temperature until experimentation in a shuttlebox. Total sample size (n), avoidance temperatures (temperature when a passage between chambers occurs), distance travelled, total number of passages, and length of weight of each experiment are shown. All values are mean±SD, with different letters denoting significant differences between groups.

| Species (age) | Incubation Treatment | n | Upper Avoidance (°C) | Lower Avoidance (°C) | Distance (m) | Number of Passages | Length (mm) | Weight (g) |
| --- | --- | --- | --- | --- | --- | --- | --- | --- |
| Lake whitefish (8 months) | 2°C | 31 | 17.54<br>(± 0.73) | 15.41<br>(± 0.81) | 193<br>(± 47) | 183<br>(± 113) | 55<br>(± 5) | 1.13<br>(± 0.32) |
|  | 5°C | 29 | 17.23<br>(± 1.09) | 14.93<br>(± 1.38) | 173<br>(± 67) | 146<br>(± 121) | 56<br>(± 6) | 1.25<br>(± 0.39) |
|  | 8°C | 25 | 17.74<br>(± 0.52) | 15.63<br>(± 0.57) | 191<br>(± 60) | 170<br>(± 124) | 55<br>(± 9) | 1.18<br>(± 0.51) |
| Lake whitefish (12 months) | 2°C | 31 | 17.14<br>(± 2.45) | 14.01<br>(± 2.62) | 247<br>(± 112) | 51<br>(± 95) | 114<br>(± 12) | 10.59<br>(± 3.04) |
|  | 5°C | 40 | 17.72<br>(± 2.26) | 14.68<br>(± 1.94) | 196<br>(± 71) | 51<br>(± 65) | 114<br>(± 11) | 10.76<br>(± 3.59) |
|  | 8°C | 32 | 17.41<br>(± 1.85) | 14.24<br>(± 2.16) | 209<br>(± 97) | 31<br>(± 55) | 114<br>(± 8) | 10.28<br>(± 2.43) |
| Round whitefish (12 months) | 0.5°C | 27 | 15.80<br>(± 0.75)<sup>a</sup> | 13.73<br>(± 0.75)<sup>a</sup> | 217<br>(± 44) | 212<br>(± 81) | 62<br>(± 5)<sup>a</sup> | 1.60<br>(± 0.45)<sup>a</sup> |
|  | 2°C | 31 | 15.29<br>(± 0.93)<sup>b</sup> | 13.18<br>(± 1.01)<sup>ab</sup> | 189<br>(± 65) | 188<br>(± 105) | 55<br>(± 6)<sup>b</sup> | *1.19<br>(± 0.35)<sup>b</sup> |
|  | 6°C | 23 | 15.11<br>(± 0.90)<sup>b</sup> | 13.04<br>(± 0.79)<sup>b</sup> | 198<br>(± 41) | 204<br>(± 79) | 62<br>(± 5)<sup>a</sup> | 1.71<br>(± 0.52)<sup>a</sup> |

**Table 2.** Temperature preference (T_pref_) of juvenile lake whitefish of different ages. Age is provided in years (yrs) and months. Age in months was estimated for previously published studies by assuming median hatch occurs within March – April, as suggested by testing dates for 1-year old fish. T_pref_ from the present study is reported from 2°C treatment groups only. Edsall (1999) used simulated lake water temperature during embryonic incubation. All other T_Pref_ data provided by Edsall (1999). Holding temperature refers to water temperature in home tanks from hatch until testing.

| Age (yrs) | Age (months) | Size (g) | $T_{pref}$ (°C) | Holding Temperature (°C) | Source |
| --- | --- | --- | --- | --- | --- |
| 0 | 4-5<sup>a</sup> | 2.8 | 15.9 | 8-11 | Edsall (1999) |
| 0 | 5-6 | 1.1-1.7 | 17-18<sup>b</sup> | -<sup>c</sup> | Opuszynski (1974) |
| 0 | 5-6 | 1.9 | 16.8 | 8-11 | Edsall (1999) |
| 0 | 8 | 1.13 | 16.04 | 15 | Present Study |
| 1 | 12 | 10.59 | 15.27 | 15 | Present Study |
| 1 | 12-13<sup>a</sup> | 3.9 | 15.6 | 8-11 | Edsall (1999) |
| 1 | 12-13 | 5.7 | 10 | -<sup>c</sup> | Opuszynski (1974) |

<sup>a</sup> Repeated measure on same cohort of fish.
<sup>b</sup> Temperature preference estimated via inspection by Opuszynski (1974).
<sup>c</sup> Information not available.

At 12 months of age, LWF from all treatment groups (2°C, 5°C, 8°C) displayed similar T_pref_ (Fig. 2B; One-way ANOVA; F _[2,100]_ = 0.0765; p=0.468). Upper and lower avoidance temperatures were comparable between all treatment groups, suggesting 12-month-old LWF were avoiding temperatures below 14.3°C and above 17.4°C on average (Table 1). Average total length and body weight were similar across all treatment groups, varying less than 1mm or 0.5g, respectively. Linear models were fit, including body weight and total length as fixed effects, to determine if there was a relationship between size and T_pref_. Model results indicated that body weight (p = 0.0678) and total length (p = 0.0607) did not significantly affect T_pref_ at 12-months-old.

Temperature exposure during rearing effects the T_Pref_ of 12-month juvenile RWF (Fig. 2C; One-way ANOVA, F _[2,78]_ =5.509; p=0.0058). The T_Pref_ RWF juveniles incubated at 2°C (13.53 ± 1.14°C) and 6°C (13.39 ± 0.99°C) as embryos displayed significantly lower T_pref_ compared to those incubated at 0.5°C (14.27 ± 0.95 °C; p=0.0216 and p=0.01, respectively), with no differences between the 2°C and 6°C groups (p=0.8764). This change in preference is reflected in recordings of upper and lower avoidance temperatures. Fish exposed to 0.5°C (15.8 ± 0.75 °C) exhibit a higher upper avoidance temperature (One-way ANOVA; F _[2,78]_ =3.51; p=0.0347; Table 1), with both 2 (15.29 ± 0.93 °C) and 6 °C (15.11 ± 0.9 °C) exposed RWF significantly decreased in comparison (p=0.0216 and p=0.01, respectively). Lower avoidance temperatures exhibit a similar trend (One-way ANOVA; F _[2,78]_ = 3.676; p=0.0298; Table 1), with 0.5°C (13.73 ± 0.75°C) treated RWF have lower avoidance temperatures than 2 (13.18 ± 1.01°C) and 6°C (13.04 ± 0.79°C) treated fish, differing significantly when compared to 6°C fish (p=0.03). Total distance travelled (One-way ANOVA, F _[2,78]_ = 1.885, p = 0.159) and number of passages (One-way ANOVA, F _[2,78]_ = 0.522, p = 0.596) were statistically similar between all treatment groups. Total length was not consistent between treatment groups (One-way ANOVA, F _[2,78]_ = 15.097, p < 0.0001) as juveniles in the 2°C group were significantly smaller than those in the 0.5°C and 6°C treatment groups (p< 0.0001). However, total length was not significantly different between 0.5°C and 6°C treatments (p=0.643). Body weight followed the same trend as total length (One-way ANOVA, F _[2,78]_ = 11.374, p = .000045, Table 1), with the 2°C group significantly smaller in body weight on average than the 0.5°C and 6°C groups (p=0.0017 and 0.0001, respectively).

As LWF and RWF reside in similar habitats, we sought to assess whether species specific differences existed in T_Pref_. LWF reared in a similar condition to RWF (2°C), exhibit an increased preference for warmer waters at 12 months of age (Fig. 3; T-test, p<0.0001). To investigate the effect of age on T_pref_ we compared 8-month-old and 12-month-old LWF incubated at the standard temperature of 2°C. Average T_pref_ for 8-month-old LWF was 16.04 ± 1.14°C compared to 15.27 ± 2.67°C for 12-month-old LWF (Fig. 4), which were not statistically different (T-test, p = 0.147).

**Figure 3.**
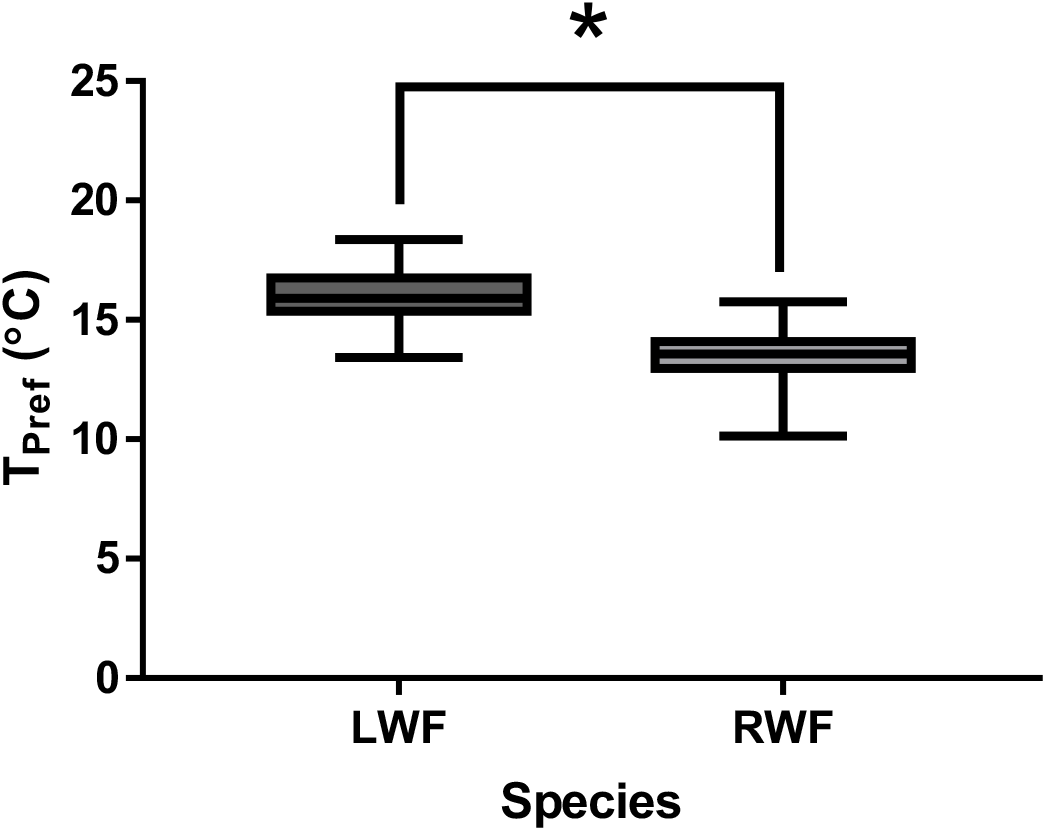
Species exhibit unique thermal preferences following rearing at a common temperature. Boxplots comparing thermal preference (T_Pref_) of 12-month-old lake whitefish (LWF) and 12-month-old round whitefish (RWF), after experiencing 2°C during embryogenesis. Lower and upper box boundaries are 25^th^ and 75^th^ percentiles of the data, with the line inside the box representing the median, and error lines encompassing the entirety of the spread of data. The symbol * is used to denote a significant difference between groups.

**Figure 4.**
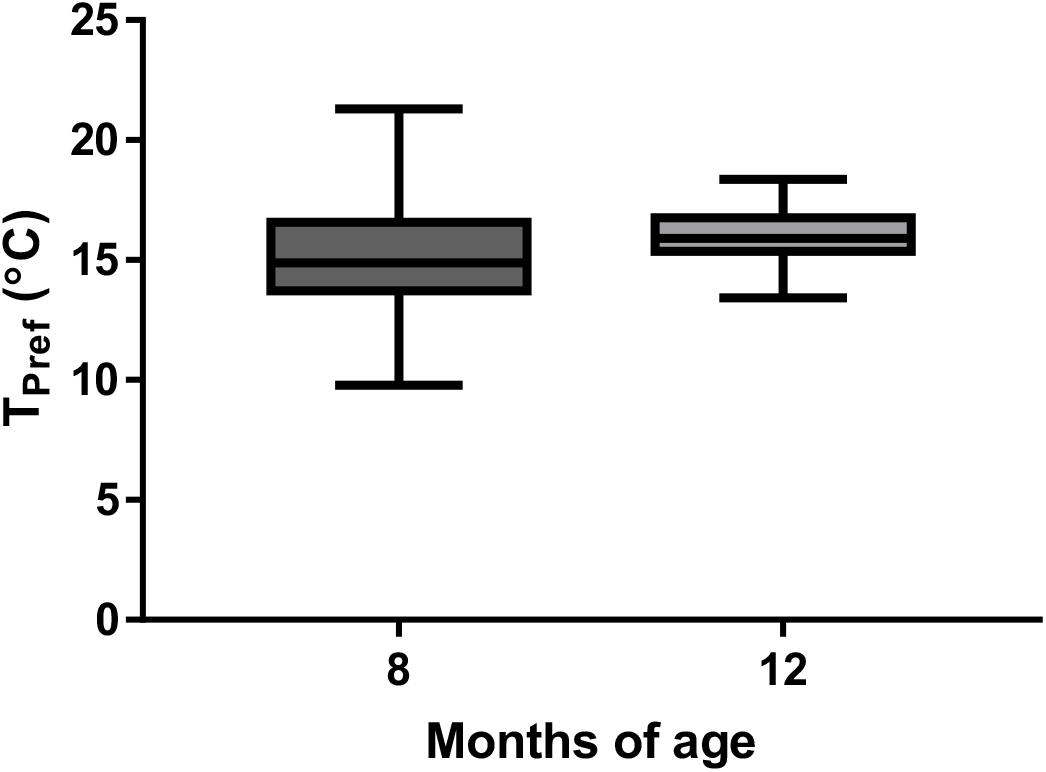
Thermal preference does not change during the juvenile stage of lake whitefish. Boxplots comparing thermal preference (T_Pref_) of 8-month-old and 12-month-old lake whitefish (LWF), after experiencing 2°C during embryogenesis. Lower and upper box boundaries are 25^th^ and 75^th^ percentiles of the data, with the line inside the box representing the median, and error lines encompassing the entirety of the spread of data.

## Discussion

Our results reveal that elevated temperatures during rearing lead to long-term changes to the T_Pref_ of RWF, but not LWF. Early-life thermal history has been shown to modulate several plastic traits, including behavior, social skills, and endocrine responses in calves (Dado-Senn et al., 2022), pigs (Johnson et al., 2018) and in fish (Jonsson and Jonsson, 2019, Li et al., 2021). While several studies have linked acute or continuous changes in temperature as the etiology of behavioral and growth changes in fish (Bartolini et al., 2015; Gillooly and Dodson, 2000; Nowicki et al., 2012; Nytrø et al., 2014; Sun and Chen, 2014), studies linking solely early-life incubation temperature to long-term perturbations or functional limitations are sparse (Jonsson and Jonsson, 2019; Scott and Johnston, 2012). Here, we provide evidence that the resulting preference of temperature for fish at the juvenile stage can be programmed during embryogenesis, but this phenotypic plasticity is species specific, and the mechanisms behind these changes are abstruse.

Most teleost fish function as ectotherms, requiring a conserved and coordinated suite of physiological and behavioral responses to navigate changes in ambient temperature (Haesemeyer, 2020; Stevens and Fry, 1970). For example, adults can preferentially swim to more optimal habitats when encountering thermal stress, such as effluent from power plants (Neill and Magnuson, 1974). These behavioral responses are key, allowing animals to maintain their metabolic optimums (Haesemeyer, 2020). At the embryonic stage, fish would be subjected to environmental temperatures without a recourse to navigate to more appropriate conditions. Several studies have described impacts of elevated incubation temperature during embryogenesis, with clear delineations of reductions in survivability and growth, but also in perturbed metabolism (Barrionuevo and Burggren, 1999; Bozek et al., 1990; Marty et al., 1990). In response to direct increases in ambient temperature, there is a positive linkage to metabolism (Fry and Hochachka, 1970). LWF embryos incubated at constant elevated temperatures display increased oxygen consumption (Eme et al., 2015), but it is unknown whether this difference in metabolism persists to the juvenile stage in this species. Other studies have reported increases in post-embryonic metabolism following embryonic incubations in elevated water temperatures, such as in the razorback sucker (*Xyrauchen texanus;* (Bozek et al., 1990), zebrafish (*Danio rerio;* Barrionuevo & Burggren, 1999), Arctic charr (Salvenlinus alpinus; (Huuskonen et al., 2003) and Japanese medaka (*Oryzias latipes;* Marty et al., 2010). This suggests that the previously noted increase in metabolism (Eme et al., 2015) may persist to later life-stages. This is an important point, as a functional link between basal metabolic rate and thermal preference has been established in the common minnow (*Phoxinus phoxinus*), demonstrating that fish with higher metabolic rates may prefer colder temperatures as juveniles (Killen, 2014). While this could then suggest that RWF reared at 0.5°C have a lowered metabolism, future studies would be required to confirm whether higher metabolism is at the root of lower T_Pref_ in RWF reared at 2 and 6°C.

Round whitefish appear to be more sensitive to elevations in rearing temperature than LWF. We originally hypothesized that increased temperature during incubation would lead to alterations in T_Pref_, based upon previous observations that elevated ambient temperature increases mortality in these species (Brooke, 1975; Mitz et al., 2019; Eme et al., 2018; Mueller, 2017; Lee et al., 2016; Lim et al., 2017; Lim et al., 2018). At 2°C, RWF experience nearly 30% increase in mortality compared to LWF, with no embryos surviving continuous exposure to 8°C (Lim et al., 2018). RWF appear to exhibit considerable sensitivity to thermal challenges, forming our rationale to reduce the thermal regime RWF were exposed to in this study (6°C), and the inclusion of the 0.5°C incubation group. Serendipitously, this reduction revealed that 2 and 6°C appear to be capable of imparting long-term alterations to T_Pref_ behavior. This possibly presents an advantage in natural settings to whitefish reared in colder water, as fish experiencing a lower temperature during embryogenesis would then prefer a higher ambient temperature at the juvenile stage. A higher temperature is typically found higher in the water column and may present a greater food supply. Indeed, zooplankton, a major food source for larval and juvenile whitefish (Freeberg et al., 1990), is commonly found in higher abundances at warmer and shallower water (Berger et al., 2006). Taken together, rearing in colder water might provide slight behavioral advantages for RWF, suggesting that ever-increasing ambient temperature driven by anthropogenic practices may be detrimental for this species.

Apart from temperature stressors, RWF appear to be more sensitive to environmental perturbations than LWF. Population declines have been observed in RWF in New York State (Bouton and Stegemann, 1993; Conley et al., 2021), leading to these fish being labelled vulnerable in this state (Bouton and Stegemann, 1993). Comparing RWF to LWF, the former has historically had a smaller distribution than LWF in North America, with LWF distribution extending farther south beyond the great lakes (Ebener et al., 2008). While the specific causes of these declines are unknown, others have speculated this could be explained by the general sensitivity of RWF to abiotic stressors. For example, acid rain has impacted the Adirondack lakes of New York State, lowering pH, and increasing aluminum and mercury, which interfere with reproduction and survival in these fish (Conley et al., 2021). Moreover, exposure to morpholine, a chemical used to prevent corrosion and damage to water pipes and is used as an additive in fossil fuels, leads to increased mortality and reduced body size in RWF, when compared to LWF at supraenvironmental levels (Lim et al., 2018; Thome et al., 2016). In this study, comparisons between LWF and RWF revealed that when raised at a similar temperature, LWF exhibit a higher T_Pref_ than RWF (Fig. 2). As adults, LWF occupy deeper (18-90 m) limnetic water, with RWF residing in shallow littoral depths (Bailey, 1964; Cucin and Regier, 1965; Rawson, 1951). However, as larvae and juveniles, round and lake whitefish are found feeding along shorelines in shallow water, before gradually moving to deeper waters (Faber, 1970; Hogman, 1971). The difference in preferred temperature may support the observation of these species overlapping, but occupying distinct niches and resources (Eberts et al., 2016).

We originally suspected that changes in size and age may play a role in determining T_Pref_ of whitefish. Previous studies have shown a significant relationship between these variables and preference of temperature in LWF, with T_Pref_ decreasing as the animal grew/age (Edsall, 1999; Opuszynski, 1974). In this study, we performed assessments to investigate both factors using LWF, performing a regression for T_Pref_ compared to size (weight and length), and directly comparing the 8-, and 12-month-old age class exposed to a similar ambient temperature during embryogenesis (Fig. 3). While we note no correlations of T_Pref_ with either size or age, we must acknowledge substantial differences of our study design with previous studies investigating this species. Life-stage plays a significant role in determining the preference of the animal, as their natural history dictates a transition to deeper waters as the animal ages (Hogman, 1971). In the work by Edsall (1999) and Opuszynski (1974), thermal preference was ascertained by using considerably younger and smaller LWF. This is a key point, as their comparisons were carried out using fish separated by approximately 6-7 months (Edsall, 1999; Opuszynski, 1974), a larger difference than the present study with a 4-month difference in age. Inherently tied to this age difference is a difference in growth, as our study generated a ~10-fold increase in weight from 8 to 12 months of age, and the previous studies describing a more modest increase of ~2-3 fold (Opuszynski, 1974; Edsall, 1999). Changes in growth can easily be attributed to holding temperature of post-hatch fish, with our study implementing a common garden temperature of 15°C, compared to the colder holding temperature used previously (8-11°C, Eddsall, 1999). This is an important consideration, as despite a substantially larger increase in absolute size in our study, we note no differences in T_Pref_. This leads to the proposal that developmental age plays a more significant role in determining temperature preferences in LWF. Plankton tow data points to whitefish migrating from warmer coastal waters to cooler, and deeper waters at approximately 4 months of age (Loftus, 1982; Ryan et al., 2014), which may suggest that the age classes investigated in this study may have surpassed windows of overt change in T_Pref_. Another important difference between these studies is the implementation of a vertical testing chamber (Eddsall, 1999; Opuszynski, 1974), compared to the horizontal shuttlebox used here. In vertical chambers, the coldest temperature is at the bottom of the arena, which may present a behavioral factor that was not considered in our design. In novel situations, fish will exhibit more bottom dwelling type behavior (Blaser and Rosemberg, 2012), which may drive a larger preference for colder temperature. A comparison between these behavioral paradigms would be prudent for understanding how an animal’s innate response during assessment may influence thermal preferences.

There appears to be no changes in activity levels of LWF and RWF. While the shuttlebox is not purposefully built to assess levels of general swimming, movement in this assessment may be considered a gauge of the animal’s exploratory behavior to seek an optimal environment. In search of a preferred temperature, LWF and RWF of all age classes move to equivalent levels and exert a similar number of chamber transitions across all temperature treatments. Increases in temperature lead to increases in locomotion, and anxiety-like behaviours (Angiulli et al., 2020; Biro et al., 2010), with evidence suggesting that the imprinting of temperature in early-life can lead to long-term changes in behavioral responses (Li et al., 2021). While we did not explicitly assess general swimming in LWF and RWF, the results generated here suggests that early-life rearing temperature does not effectively alter behavior during thermal testing.

In conclusion, this study demonstrates a persistent effect of increased embryonic incubation temperature on the thermal preference of juvenile RWF. Benthic water temperatures of 2°C represent a winter of low ice cover but is sufficient to alter preferences in these fish. Given that temperatures are expected to increase (0.1°C/year; Austin & Colman, 2007), and thermal effluents impact coastal water temperature, these results raise concern for a fish species that has been considered in decline (Bouton and Stegemann, 1993; Conley et al., 2021), and are rarely seen in abundant amounts ecologically (Mraz, 1964). Coastal embayments provide a thermal refuge during the spring warming (Ryan and Crawford, 2014) and ice-free conditions facilitate a spring bloom of primary productivity which is important for survival of larval whitefish (Faber, 1970). Round whitefish seeking cooler water temperatures may avoid prime nursery grounds, which would put them at a disadvantage compared to other conspecifics. Cold-adapted freshwater fish are among the taxa most vulnerable to climate change but receive a fraction of the research and conservation efforts of terrestrial species (Pacifici et al., 2015). This study highlights the importance of examining sub-lethal thermal effects and thermal plasticity of cold-adapted species. Future studies seeking to understand the role of metabolism on thermal preference are prudent, as this technique provides a non-invasive assessment of environmental performance that may be used to determine at risk-populations environmentally.

## List of Symbols/Abbreviations

LWF: Lake Whitefish (*Coregonus clupeaformis*)
RWF: Round Whitefish (*Prosopium cylindraceum*)
T_pref_: Temperature Preference
°C: Degrees in Celsius
mL: Milliliter
min: Minute
mm: Millimeter
g: gram

## Acknowledgements

We would like to thank undergraduates Urvi Pajankar and Akanksha Arora for their contributions to whitefish husbandry. Tim Drew from the MNRF Sharbot Lake Whitefish Culture Station, and Dr. Fei Luo with Ecometrix for providing lake and round whitefish embryos respectively. Dr. Ben Bolker for statistical discussion, and Dr. Grant McClelland for feedback on design and analysis.

## Author Contributions

Conceptualization: A.A.H., J.Y.W., R.G.M, C.S.M.; Methodology: A.A.H., J.Y.W.; Resources: A.A.H., M.L.M.F., L.L.; Formal Analysis: A.A.H., W.A.T.; Investigation: A.A.H., H.M., B.A, M.M.L.F. L.L.; Data Curation: A.A.H., H.M., W.A.T.; Writing – original draft: A.A.H., W.A.T.; Writing – review and editing: W.A.T., A.A.H., J.Y.W., M.L.M.F., D.R.B., C.M.S., R.G.M.; Supervision: J.Y.W.; Project administration: A.A.H., J.Y.W., L.L.; Funding Acquisition: J.Y.W., D.R.B, C.M.S., R.G.M.

## Data availability

Data are available from GitHub: https://github.com/WilsonToxLab/Shuttlebox-Thermal-Preference

